# Integrative analyses of multi-tissue Hi-C and eQTL data demonstrate close spatial proximity between eQTLs and their target genes

**DOI:** 10.1101/392266

**Authors:** Jingting Yu, Ming Hu, Chun Li

## Abstract

Gene regulation is important for cells and tissues to function. At the genomic level, it has been studied from two aspects, the identification of expression quantitative trait loci (eQTLs) and identification of long-range chromatin interactions. It is important to understand their relationship, such as whether eQTLs regulate their target genes through physical chromatin interaction. Although previous studies have suggested enrichment of eQTLs in regions with a high chromatin interaction frequency, it is unclear whether this relationship is consistent across different tissues and cell lines and whether there would be any tissue-specific patterns. Here, we performed integrative analyses of eQTL and high-throughput chromatin conformation capture (Hi-C) data from 11 human primary tissue types and 2 human cell lines. We found that chromatin interaction frequency is positively correlated with the number of genes having eQTLs, and eQTLs and their target genes are more likely to fall in the same topologically associating domains than that expected from randomly generated control datasets. These results are consistent across all tissues and cell lines we evaluated. Moreover, in dorsolateral prefrontal cortex, spleen, hippocampus, pancreas and aorta, tissue-specific eQTLs are enriched in tissue-specific frequently interacting regions. These results reveal a more detailed picture of the complicated relationship between different mechanisms of gene regulation.

**Author summary:** Whole-genome gene regulation has been studied in tissues and cell lines from multiple perspectives, including identification of expression quantitative trait loci (eQTLs) and identification of long-range chromatin interactions. These two complementary approaches focus on different aspects of gene regulation, one being statistical across individuals while the other being physical within a sample. Integrating results from these two approaches will help us understand their relationships, such as whether eQTLs regulate their target genes through physical chromatin interaction. We performed comprehensive analyses using data from multiple human tissues and cell lines, and showed that chromatin interaction frequency is positively associated with eQTL results in all evaluated tissues and cell lines. The observed relationships also displayed tissue-specific pattern in some tissues. Our results revealed a more detailed picture of the complicated relationship between the different mechanisms of gene regulation.

## Introduction

Gene regulation is important for cells and tissues to function. Differences in gene regulation are often responsible for cellular and morphological differences between cell lines and tissues. The advancement of high-throughput technologies such as DNA and RNA sequencing and SNP chips allows researchers to study gene regulation at the genomic level and from multiple perspectives. On the one hand, motivated by the likely functional importance of genetic variants in gene regulation, many studies have focused on identifying expression quantitative trait loci (eQTLs), which are genetic variants statistically associated with gene expression across individuals [1–4]. eQTLs can regulate the expression of their target genes by altering *cis*-regulatory elements (CREs) such as enhancers, promoters, insulators, mediators, etc [5–7]. On the other hand, analyses of chromatin spatial organization have established the importance of chromatin interaction in gene regulation [8–10]. For example, by forming long-range chromatin interactions, CREs can regulate the expression of their target genes hundreds of kilobases (Kb) away [11–13]. High-throughput chromatin conformation capture (Hi-C) has been widely adopted to provide a genome-wide view of chromatin interactions within a tissue or cell line [14–17].

These two complementary approaches focus on different aspects of gene regulation. eQTL results are statistical across individuals and require an associated SNP, while chromatin interactions are physical within a sample and do not require a polymorphism to be present. It is desirable to integrate the results of these two approaches to better understand their relationships, such as whether eQTLs regulate their target genes through chromatin interactions. Analyzing Hi-C data from human IMR90 fibroblasts and embryonic stem cells, Duggal *et al.* [18] showed that eQTLs are spatially close to their target genes, especially for those located within the same topologically associating domains (TADs) and overlapping with CREs, and that genomic regions containing eQTLs tend to have a higher chromatin interaction frequency. Using Hi-C data generated from several human cell lines, the Genotype-Tissue Expression (GTEx) study [3] has shown that eQTLs enriched for CREs are in close spatial proximity with their target gene promoters. However, these two studies relied on eQTL results that were aggregated from different tissues and cell lines. Therefore, it is unclear if the observed relationships are consistent across different tissues or cell lines and if there would be any tissue-specific patterns in the observed relationships.

Recently, Schmitt *et al.* [19] generated Hi-C data for 14 human primary tissues and 7 human cell lines, and found that frequently interaction regions (FIREs) are enriched for CREs and display patterns that are tissue and cell-type specific. Meanwhile, the GTEx study [3] performed genome-wide mapping of eQTLs across 48 human tissues, and found that eQTLs are enriched in CREs and have varied genetic effect across tissues. There are 11 tissues and 2 cell lines that overlap between these two sources (**S1 Table** and see **S1 File**). In this paper, we analyze these data to evaluate the relationships between eQTLs and chromatin interactions across multiple tissues and cell lines.

We first performed a comprehensive evaluation of the association between eQTLs and chromatin interactions, as well as the enrichment of eQTL results in TADs. We found that chromatin interaction frequency is positively correlated with the number of genes that are significantly associated with eQTLs (i.e., eGenes), and eQTLs and their target genes are more likely to reside within the same TAD than randomly generated control datasets. The results are consistent across all tissues and cell lines we evaluated. Since both eQTLs and FIREs are known to be highly tissue-specific, their relationships may vary from tissue to tissue. Thus we characterized the association between eQTL and chromatin interaction for individual tissues, and found that tissue-specific eQTLs are enriched in tissue-specific FIREs.

To the best of our knowledge, our study is the first to reveal the relationship between eQTLs and chromatin interactions across multiple tissues, and to characterize the relationship between tissue-specific eQTLs and tissue-specific FIREs. Some relationships are consistent across tissues and cell lines, while a few others display tissue-specific patterns. These results provide a comprehensive survey of the relationship between different mechanisms of gene regulation, and may help improve our understanding of their roles in gene regulation mechanisms.

## Results

### Chromatin interaction frequency is positively correlated with the number of eGenes

If chromatin spatial organization affects how eQTLs regulate their target genes, one would expect that genomic regions mapped with eQTL-gene associations would interact frequently. To test this hypothesis, we fitted negative binomial regression models to evaluate the relationship between the number of eGenes and chromatin interaction frequency. After adjusting for genomic distance and the total number of tested genes, the number of eGenes showed significantly positive effects on chromatin interaction frequency across all tissues and cell lines (**Fig 1A**). For example, in spleen, the effect of the number of eGenes is estimated to be 0.19 (p value <2.2e-16), indicating that chromatin interaction frequency would be 1.21 (= *e*^0.19^) times higher for every extra eGene in a bin pair. The magnitude of the effects varies across tissues and cell lines, ranging from 0.05 to 0.19. However, the effects are similar for tissues from the same organ, such as the two brain tissues, DLPFC and hippocampus. Moreover, genomic distance has a significant negative effect on chromatin interaction and the effects are similar across all tissues and cell lines. This is as expected because chromatin interaction frequency between two genomic regions tends to decrease as their genomic distance increases [16]. We also stratified the data by distance and the total number of tested genes. S1 Fig contains the results for GM12878. Some strata display positive association between chromatin interaction frequency and the number of eGenes, although the overall results are noisier since stratification greatly reduced sample size.

**Figure 1.**
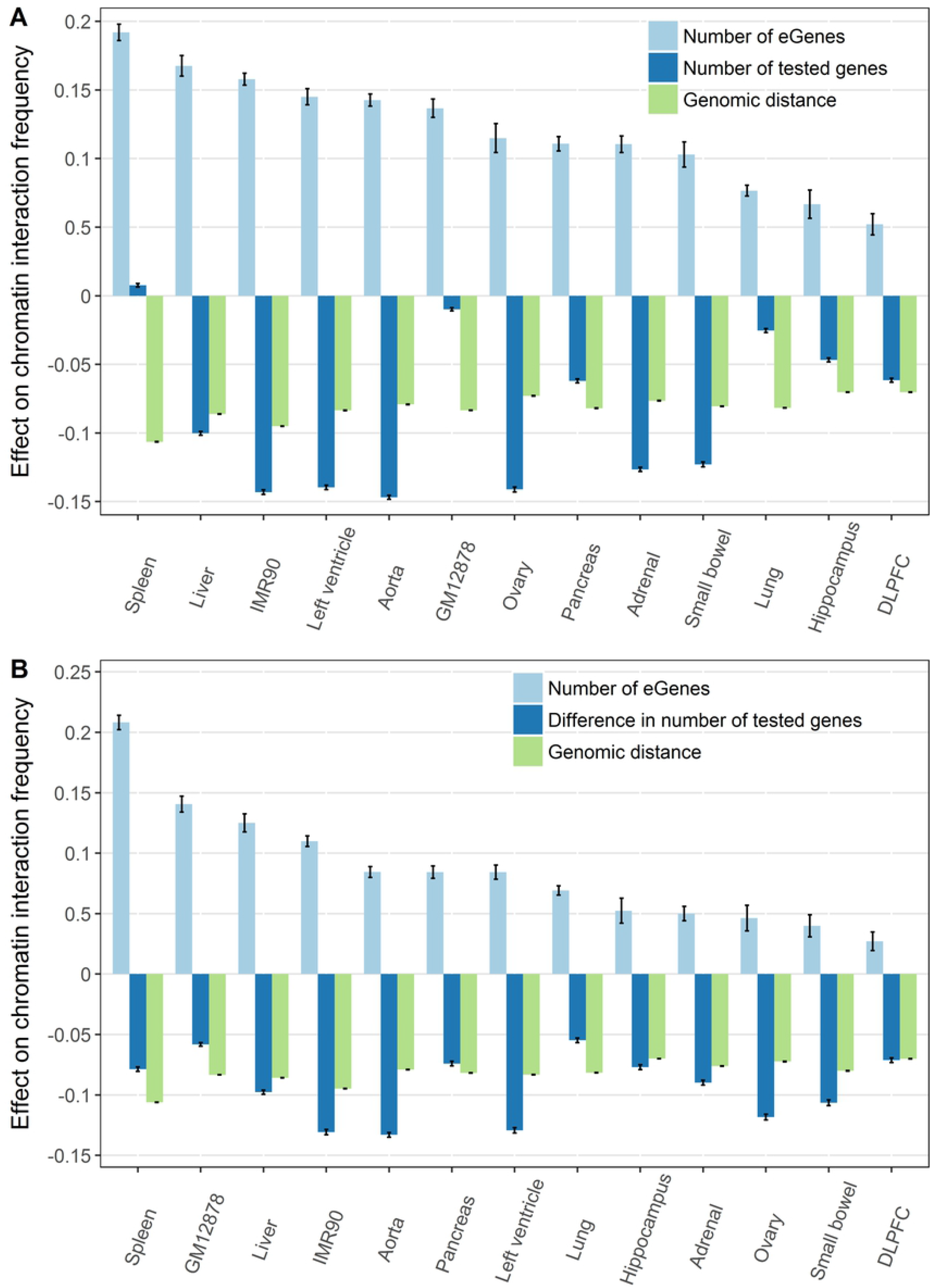
Chromatin interaction frequency is positively correlated with number of eGenes. Two negative binomial regression models were fitted for chromatin interaction frequency. **(A)** Bat plot showing the estimated effects of number of eGene (light blue), total number of tested genes (dark blue), and genomic distance (green). **(B)** Bar plot showing estimated effects of number of eGene (light blue), difference of tested genes (dark blue), and genomic distance (green).

Lieberman-Aiden *et al.* [16] have discovered the A and B compartments, which are correlated with relatively high and low gene density, respectively. We thus fitted a regression model with the absolute difference in the number of tested genes between the two genomic regions in place of the total number of tested genes (**Fig 1B**). As we expected, for all tissues and cell lines, the difference in gene density has a significant negative effect on chromatin interaction frequency, indicating that genomic regions with a larger discrepancy in gene density tend to interact less frequently. The effects of the number of eGenes are still significantly positive, although their magnitudes are less than those in the previous model except for spleen.

In addition to the two models above, we repeated the analyses with the fraction of eGenes in place of the number of eGenes (**S2 Fig**). Consistent with the results from first two models, the eGene fraction also has a significant positive effect on chromatin interaction frequency. Collectively, these results confirm that eQTL-gene associations are positively correlated with chromatin interaction frequency, which implicates the potential regulatory role of chromatin interactions in facilitating eQTLs regulating their target genes.

### The eQTL-gene associations are enriched in TADs

Since genomic regions within the same TAD are known to interact more frequently than those in different TADs [20, 21], we next examined whether eQTL-gene associations are enriched within TADs. For each tissue and cell line, we simulated a set of pseudo-SNP-gene pairs through matching the genomic distances of true eQTL-gene associations. By comparing the fraction of associations mapping in same TADs between the simulated and real data, we found that the real data showed significantly higher fraction of eQTL-gene pairs falling in same TADs than the simulated data, and this is true across all tissues and cell lines. For example, 74% of real eQTL-gene associations and 70% of simulated pairs in GM12878 were inside TADs (Fisher’s exact test, p-value < 2.2e-16). We further stratified the data by distance between the eQTL and its associated gene. We found that majority of eQTL-gene associations at distance from 40Kb to 400Kb are significantly enriched in TADs (**Fig 2** and **S3 Fig**). For example, for GM12878, the real eQTL-gene pairs had a significantly higher fraction inside TADs than the simulated data at distance 40Kb to 280Kb (**Fig 2A**).

**Figure 2.**
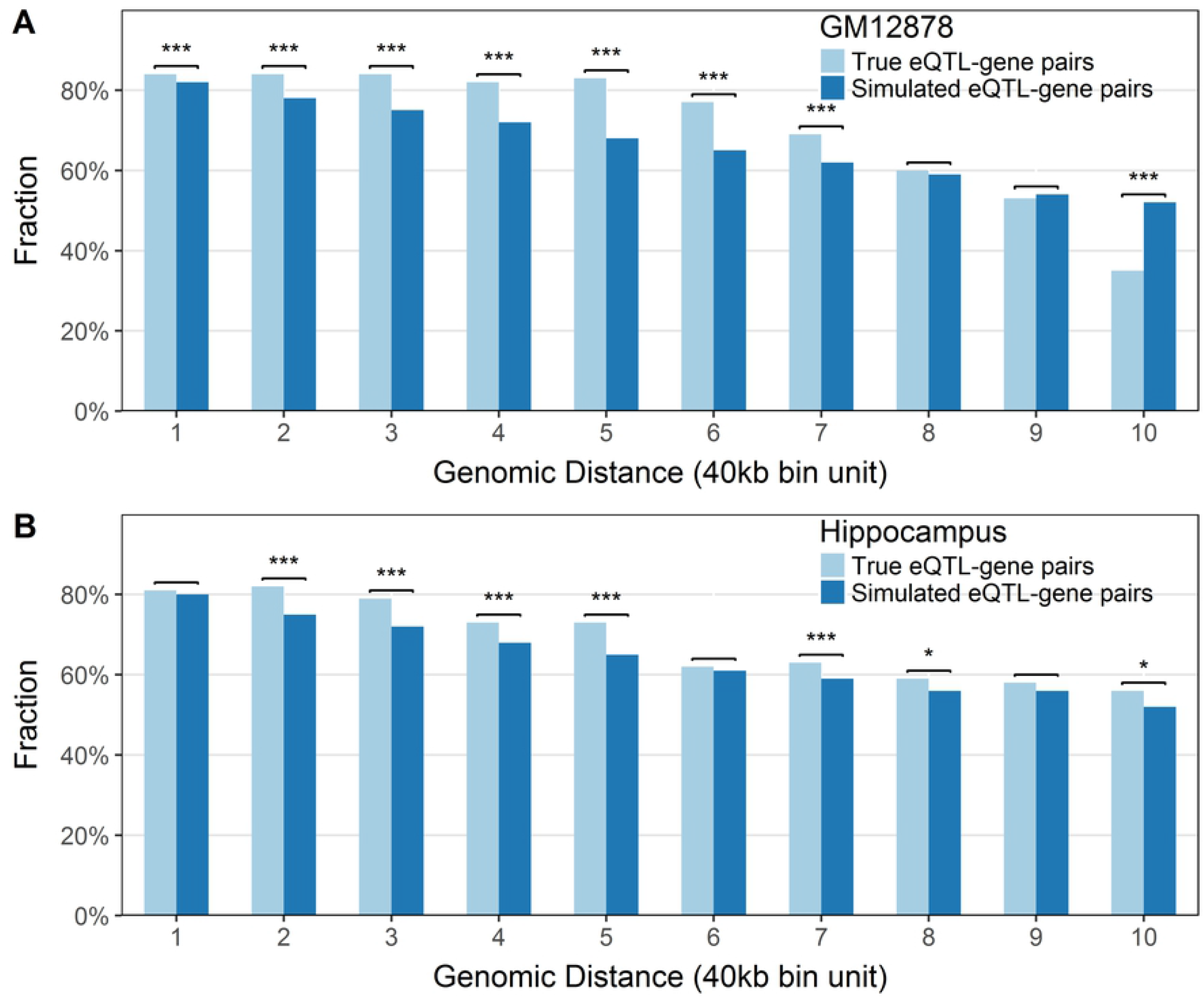
The eQTL-gene associations are enriched in TADs. **(A)** Bar plot for GM12878 showing the fractions of eQTL-gene pairs that mapped inside TADs (y-axis) from the true (light blue) and simulated data (dark blue). All the pairs are grouped by the distances between the eQTLs and their associated gene, which are showed as unit of 40Kb in x-axis. Results are showed for pairs with distance within 400Kb. At every fixed distance, fractions from real and simulated distance are compared and the significance of difference resulted from the Fisher’s exact test are mark by stars on the top. ‘***’ means p-value <0.001 and ‘*’ means p-value <0.05. **(B)** Same as A, except for hippocampus tissue (The p-values are in **S2 Table**).

### Tissue-specific eQTLs are enriched in tissues-specific FIREs

The GTEx project has identified many tissue-specific eQTLs, with effects in only one or a few tissues [3]. Meanwhile, FIREs identified from Hi-C data also showed strong tissue specificity [19]. Intrigued by the high tissue specificity of both eQTLs and FIREs, we next asked whether the association between eQTL results and Hi-C data is also tissue-specific. Specifically, we first identified tissue-specific FIREs and tissue-specific eQTLs based on the data from Schmitt *et al.* and the GTEx project, respectively. For the 11 tissues we considered, a total of 349,311 eQTLs were tissue-specific. By design, all eQTLs detected by the GTEx study are within 1Mb of the TSS of tested genes. On average, 3,488 FIREs were identified per tissue (**S1 Table**) and 18% of them were tissue-specific. We found a significant enrichment for tissue-specific FIREs in regions near the TSS of genes tested in GTEx. As showed in **Fig 3**, for each tissue, we observed a significantly higher fraction of tissue-specific FIREs within 1Mb of the TSS of tested genes than in the rest of the genome. These results are consistent with those reported in Schmitt *et al.* [19], which have showed that tissue-specific FIREs are near tissue-specifically expressed genes. Our results demonstrate that a similar relationship holds when the FIREs and gene expression data are from different sources and matched simply by tissue name.

**Figure 3.**
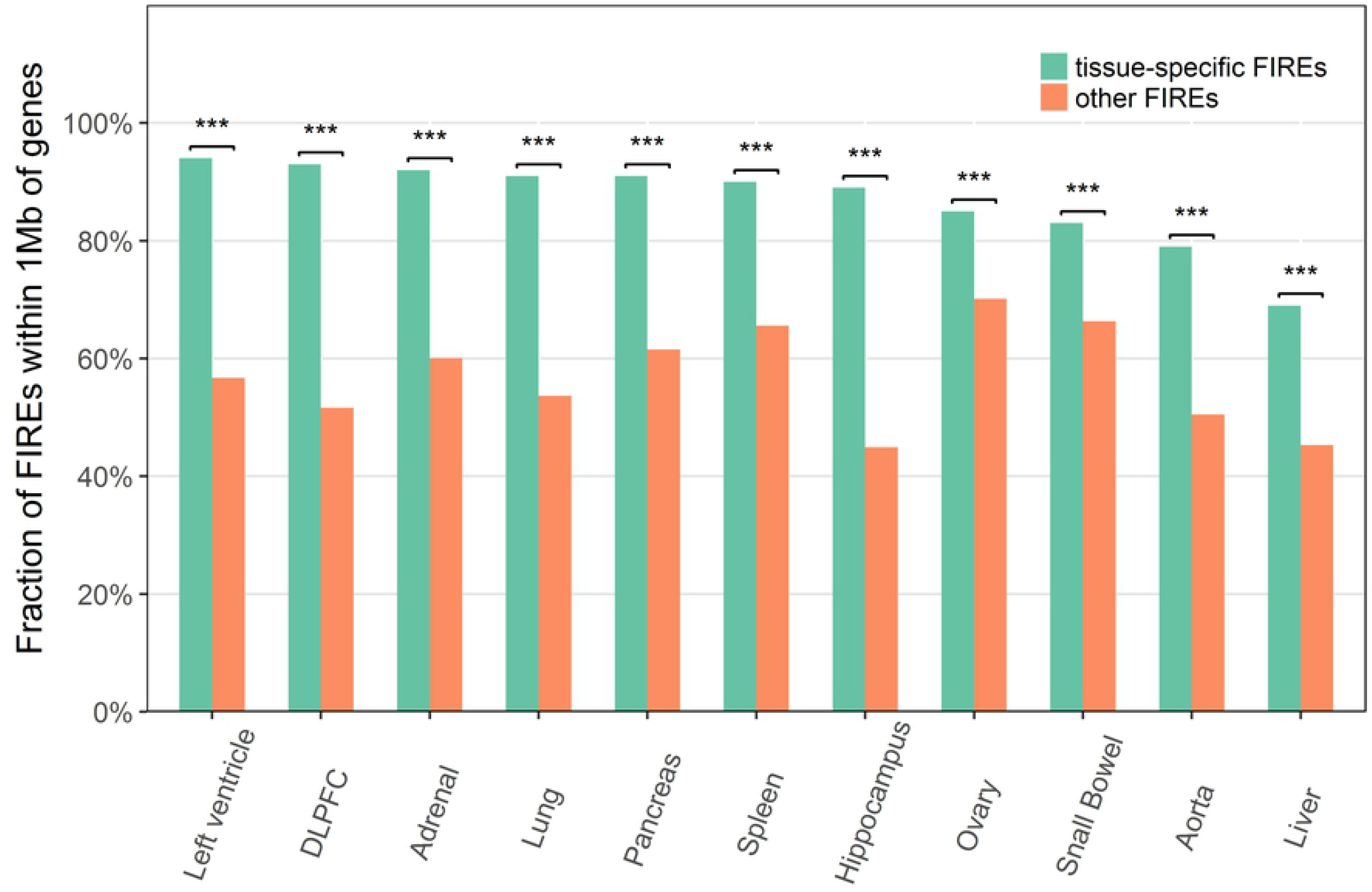
Tissue-specific FIREs are enriched near the genes tested in GTEx. Bar plot showing the fractions of FIREs (y-axis) that located within 1Mb of tested genes in GTEx for tissue-specific FIREs (green) and the rest of FIREs (orange) across tissues (x-axis). For each tissue, we evaluated the enrichment of tissue-specific FIREs within 1Mb of genes by the Fisher’s exact test. The significance of enrichment are showed as star on the top of bars. ‘***’ means p-value <0.001 (The p-values are in **S3 Table**)

We then examined whether tissue-specific eQTLs are enriched in tissue-specific FIREs. Since all the eQTLs are within 1Mb of the TSS of tested genes by the design of the GTEx study, we focused on FIREs that are also within 1Mb of the genes tested in GTEx for the tissue of interest. For each tissue, we compared the fraction of tissue-specific eQTLs mapped to tissues-specific FIREs and to other FIREs. Among the 11 tissues we evaluated, five of them (DLPFC, spleen, hippocampus, pancreas and aorta) have significant positive association after Bonferroni correction (**Fig 4**). In these five tissues, tissue-specific eQTLs are enriched in tissue-specific FIREs, suggesting that there may be synergy between eQTLs and chromatin spatial organization for gene regulation. However, a negative association was found in lung, suggesting a more complicated relationship between eQTLs and chromatin spatial organization. The results for other tissues were not significant. We also repeated the analysis for the cell lines GM12878 and IMR90, and obtained are not significant results (Fig 4). Taken together, these results demonstrate strong tissue-specificity in the association between eQTLs and FIREs for some tissues.

**Figure 4.**
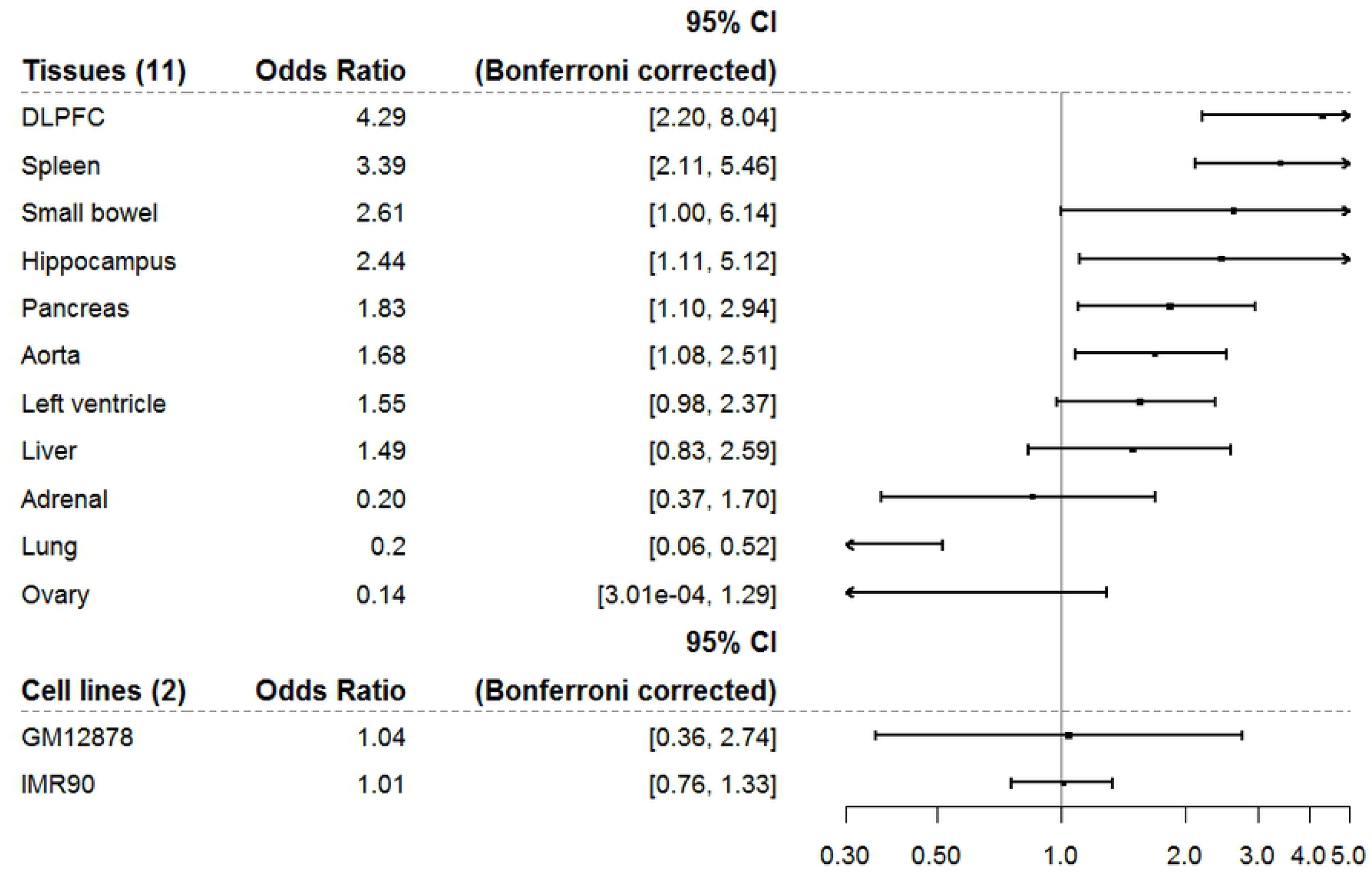
Tissue-specific eQTLs are enriched in tissues-specific FIREs. Forest plot showing the enrichment of tissue-specific eQTLs at tissue-specific FIREs for 11 tissues and 2 cell lines. We focused on the regions within 1Mb of tested gene in GTEx and tallied the number of eQTLs according to whether they are tissue-specific eQTLs and whether they fall in tissue-specific FIREs. For each tissue/cell line, we performed chi-square test to evaluate the association between tissue-specific eQTLs and tissues-specific FIREs. The odds ratios and 95% confidence intervals (after Bonferroni correction) are showed in the plot. The scale of odds ratio is in log-scale.

## Discussion

Chromatin spatial organization and eQTLs are known to be involved in gene regulation. In this work, we systematically studied the relationship between eQTL-gene association and chromatin interaction across 11 tissues and 2 cell lines. To the best of our knowledge, this is the most comprehensive study on this topic up to date. We found that chromatin interaction frequency is positively correlated with the number of eGenes in all tissues and cell lines we evaluated. Moreover, we found that eQTL-gene associations are enriched in TADs. Since both eQTLs and FIREs are known to be tissue-specific, we further evaluated the tissue-specificity of the relationship between eQTL-gene associations and chromatin interactions. We found that in DLPFC, spleen, hippocampus, pancreas, and aorta, tissue-specific eQTLs are significantly enriched in tissue-specific FIREs. This results highlight the tissue-specific manner of the positive relationship between eQTLs and chromatin interactions. However, lung showed a significant negative association between tissue-specific eQTLs and tissue-specific FIREs, which might be due to more complicated mechanisms or high tissue heterogeneity. Our data demonstrate the complexity of the relationship between eQTL-gene associations and chromatin interactions.

The tissue-specificity of the relationship between eQTLs and chromatin interactions can be useful for identifying tissue-specific genes that are likely to be regulated by eQTLs through chromatin interactions. For example, in the brain cortex tissue DLPFC, there are 2,954 tissue-specific eQTLs identified from the GTEx data and 323 tissue-specific FIREs identified from the Hi-C data. When both factors are considered, we identified 32 DLPFC-specific eQTLs located in the tissue-specific FIREs. These eQTLs are significantly associated with 4 genes, including ADGRB2 (adhesion G protein-coupled receptor B2), WASF3 (WAS protein family member 3), SPEF2 (sperm flagellar 2), and XPA (xeroderma pigmentosum complementation group A). Among these genes, ADGRB2, which encodes a transmembrane signaling receptor [22], has a brain-specific developmental expression pattern and its expression level is increased as the development of the brain progressed [22]. The TSS of the ADGRB2 gene (chr1:32,192,718) is ~47Kb from a brain-specific FIRE (chr1:32,240,000-32,320,000).

We matched the chromatin interaction data to the eQTL data simply by tissue name. For each tissue, Hi-C data from Schmitt *et al.* were from a single tissue donor that is different from the samples used in the GTEx study. In addition, there might also be a mismatch in what part of the tissue the samples came from. These factors might have introduced noises in our analyses. Some of our statistically significant results might be stronger if the data had been generated from the same source of samples with well controlled tissue heterogeneity.

The eQTL results used in our study were available only for SNP-gene pairs that are within 1Mb distance. The power of the eQTL analysis is largely determined by sample size. Because of these issues, we might have missed some SNP-gene associations in our analyses. In addition, our TAD enrichment analysis did not account for linkage disequilibrium (LD) between eQTLs. The effects of LD, if any, are probably canceled out between the real and simulated datasets.

In summary, we have performed a comprehensive study to reveal the significant positive correlation between eQTL results and chromatin interactions across multiple tissues and cell lines, and showed that for several tissues this correlation is tissue-specific. Our results provide insights into the functional roles of chromatin spatial organization and eQTLs in gene regulation mechanisms.

## Materials and methods

### Data description

In our analysis, we used Hi-C and eQTL data of 11 primary human tissues and 2 cell lines from Schmitt *et al.* [19] and GTEx project [3], including the lymphoblastoid cell line GM12878, the fetal lung fibroblast cell line IMR90, and adrenal, aorta, dorsolateral prefrontal cortex (DLPFC), hippocampus, left ventricle, liver, lung, ovary, pancreas, small bowel and spleen tissues (**S1 Table** and see **S1 File**). In all our analyses, we focus on the autosomes. The reference genome is hg19.

The Hi-C data contained over 2.9 billion raw intra-chromosomal unique paired-end reads on 13 samples in total, out of which >1 billion are long-range read pairs (>15Kb). An average of 2,068 TADs per sample were identified in the original study [19].

### Regression analysis of chromatin interaction frequency

We first evaluated the relationship between eQTL results and chromatin interaction frequency on the 11 tissues and 2 cell lines using regression analysis (see **S1 File**). Specifically, we considered autosomal chromosomes in 40kb bin resolution, and for every bin pair (*i, j*), we defined the following features: 1) chromatin interaction frequency (*I_Hi-c_*), 2) the number of tested genes with transcription start site (TSS) mapped to bin *i* (*G_i_*), 3) the number of tested genes with TSS mapped to bin *j* (*G_j_*), 4) the total number of tested genes (*G_T_* = *G_i_* + *G_j_*), 5) the total number of eGenes (*G_s_*), and 6) the genomic distance between bin *i* and bin *j* (*D* = |*i – j*|). If a tested gene or an eGene is in bin *i* (or *j*), its corresponding SNP or eQTL must be in bin *j* (or *i*). We focused on genomic regions that are known to be involved in both data sources (Hi-C and GTEx). Specifically, for every tissue or cell line, we considered bin pairs that contain at least one tested SNP-gene pair in which the tested gene has fragments per kilobase of transcript per million mapped reads (FPKM)>1 in the corresponding tissue or cell line in the Schmitt *et al.* study [19].

We first performed negative binomial regression of chromatin interaction frequency on the number of significant genes. Since chromatin interaction frequency is known to be affected by genomic distance and gene density [16], we included distance and the total number of tested genes as covariates in our model:

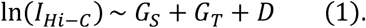

We also considered an alternative model by replacing *G_s_* with the fraction of significant genes:

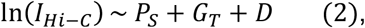

where 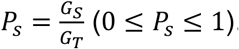. In addition, we also conducted stratified analyses over subsets stratified by the total number of tested genes and the distance between bins.

Previous studies have shown that chromatin interactions are less frequent between gene-dense A compartments and gene-poor B compartments than those within the same compartments [16]. We thus considered if the unevenness in the distribution of tested genes between two bins in a bin pair could affect chromatin interaction frequency. Specifically, we replaced *G_T_* by *D_T_* = |*G_i_ – G_j_*| and repeated the above analyses:

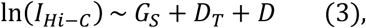

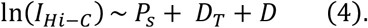

### Enrichment analysis of eQTL-gene association in TAD

We next evaluated if eQTL-gene associations are enriched in TAD. For each tissue or cell line, as a control, we simulated a set of SNP-gene pairs that have the same distribution of genomic distance as the real data, and then compared the fraction of eQTL-gene pairs falling into the same TAD with that of the simulated data. Specifically, for each tissue or cell line, we randomly selected 10,000 eQTL-gene associations from the real data and obtained the linear genomic distance (*d_i_*) between the eQTL and the TSS of its associated gene. If the eQTL is upstream of the TSS, *d_i_* is positive; otherwise *d_i_* is negative. For each of these 10,000 associations, we randomly selected an autosomal gene from the list of genes tested in GTEx for the tissue or cell line, obtained its TSS position t¿, and designated *t_i_* + *d_i_* as the position of a simulated SNP as long as the position is within the chromosome.

Next, we performed the Fisher’s exact test on the 2x2 tables of the counts of SNP-gene pairs according to whether the pair is real or simulated and whether it is in the same TAD, and then repeated the analysis by stratifying the data by genomic distance ranging from 40Kb to 1 Mb.

### Identification of tissue-specific FIREs and tissue-specific eQTLs

For each of the 11 tissues, we defined tissue-specific FIREs as those detected only for that tissue and not for any of the other 10 tissues. Tissue-specific eQTLs were similarly defined using the GTEx meta-analysis results (see **S1 File**). Cell line-specific FIREs and eQTLs were similarly defined using all 13 samples we considered. For example, GM12878-specific FIREs are the FIREs detected only in GM12878 and not in any of the other 12 samples (11 tissues and IMR90 cell line).

### Enrichment analysis of tissue-specific FIREs

For each tissue, we evaluated whether tissue-specific FIREs tend to be close to genes. We counted the number of FIREs by whether the FIRE is within 1Mb of the TSS of genes tested in GTEx for the tissue and by whether it is tissue-specific, and performed the Fisher’s exact test on the 2x2 table to assess the statistical significance of the enrichment of tissue-specific FIREs within 1Mb of genes.

### Tissue-specific analysis of eQTLs and FIREs

To explore the tissue specificity of the relationship between eQTLs and FIREs, we examined whether tissue-specific eQTLs are enriched in tissue-specific FIREs for each tissue and cell line. While all eQTLs identified by GTEx are within 1Mb of the TSS of genes, we focused on FIREs that are also within 1Mb of the TSS of genes tested in GTEx. For each tissue or cell line, we counted the number of eQTLs according to whether the eQTL is tissue-specific and whether it falls in a tissue-specific FIRE. We computed the odds ratios and the corresponding confidence intervals at the 95% level after Bonferroni correction.

## Supporting information

**S1 Fig. Distribution of chromatin interaction frequency in GM12878 stratified by total number of tested genes and genomic distance.** Plots A to D are showing the distribution of interaction frequency (y-axis) with total number of tested genes from 1 to 4. With each fixed number of tested genes, the chromatin interaction frequency were further stratified by genomic distance (x-axis), which showed as unit of 40Kb bin. At each fixed distance, the distribution of interaction frequency was presented as a function of number of eGenes.

**S2 Fig. Chromatin interaction frequency is positively correlated with fraction of eGenes.Two negative binomial regression models were fitted for interaction frequency.** (A) Bat plot showing the estimated effects of fraction of eGenes (light blue), total number of tested genes (dark blue), and genomic distance (green). (B) Bar plot showing estimated effects of fraction of eGenes (light blue), difference of tested genes (dark blue), and genomic distance (green).

**S3 Fig. The eQTL-gene associations are enriched in TADs.** Same as Figure 2, except for other 10 tissues (adrenal, aorta, DLPFC, lung, liver, left ventricle, ovary, pancreas, small bowel and spleen) and 1 cell line (IMR90).

**S1 Table. Data summary for tissues and cell lines.**

**S2 Table. The p-values for tissues in Figure 2**.

**S3 Table. The p-values for tissues in Figure 3.**

**S1 File. Supplementary materials and methods.**

